# Uncovering the genetic diversity in *Aedes aegypti* insecticide resistance genes through global comparative genomics

**DOI:** 10.1101/2024.02.29.582728

**Authors:** Anton Spadar, Emma Collins, Louisa A. Messenger, Taane G. Clark, Susana Campino

## Abstract

Insecticides are essential to control the transmission of vector-borne diseases to humans and animals, but their efficacy is being threatened by the spread of resistance across multiple medically important mosquito species. An example of this is *Aedes aegypti* - a major vector of arboviruses, including Zika, dengue, yellow fever, West Nile, and Chikungunya, with widespread insecticide resistance reported in the Americas and Asia, while data from Africa is more limited. Here we investigate the global genetic diversity in four insecticide resistance associated genes: *ace-1*, *GSTe2*, *rdl* and *vgsc.* Apart from *vgsc*, the other genes have been less investigated in *Ae. aegypti*, and limited genetic diversity information is available. We explore a large whole-genome sequencing dataset of 729 *Ae. aegypti* across 15 countries including nine in Africa. Among the four genes, we identified 1,829 genetic variants including 474 non-synonymous substitutions, as well as putative copy number variations in *GSTe2* and *vgsc*. Among these are many previously documented insecticide resistance mutations which were present at different frequencies and combinations depending on origin of samples. Global insecticide resistance phenotypic data demonstrated variable resistance in geographic areas with resistant genotypes. These warrant further investigation to assess their functional contribution to insecticide resistant phenotypes and their potential development into genetic panels for operational surveillance. Overall, our work provides the first global catalogue and geographic distribution of known and new amino-acid mutations and duplications that can be used to guide the identification of resistance drivers in *Ae. aegypti* and thereby support monitoring efforts and strategies for vector control.

## INTRODUCTION

Mosquitoes of the genus *Aedes*, particularly *Aedes* (*Ae.*) *aegypti*, are responsible for the transmission of many arboviral diseases, including dengue, Zika, yellow fever, West Nile and Chikungunya, resulting in millions of infections globally per year with limited treatment and vaccination options (1). The geographical distribution of *Ae. aegypti* has expanded considerably in recent years, predominantly due to adaptation of this vector to urban environments, climate change and the globalization of human activities, thereby increasing the risk of resurgence and spread of arbovirus infections (2–4). Compounding the problem is the global emergence of insecticide resistance among *Ae. aegypti* and other mosquito species, which is threatening to jeopardise the operational effectiveness of vector control campaigns.

Resistance to the four most common classes of insecticides used against adult mosquitoes (carbamates, organochlorines, organophosphates, and pyrethroids) has now been documented worldwide. Resistance in many mosquito species has been associated with target site mutations, metabolic detoxification, cuticular alterations and behavioural avoidance (5,6) with a suite of alternative resistance mechanisms being revealed (7–10). Target site resistance is related to mutations in insecticide target genes, such as the voltage-gated sodium channel (*vgsc* also known as knockdown resistance; *kdr*), acetylcholinesterase-1 (*ace-1* also known as *AChE1*) and γ-aminobutyric acid (GABA) receptor (resistance to dieldrin; *rdl*). Mutations in glutathione-s-transferase epsilon two (*GSTe2*), which encodes an insecticide metabolising enzyme, have also been associated with resistance (5,11–13). The *vgsc* is a large protein that is an integral part of the insect nervous system. DDT (dichloro-diphenyl-trichloroethane) and pyrethroid insecticides interfere with the *vgsc* by prolonging the pore open state leading to insect paralysis and death (14). In the reference insect for this gene, *Musca domestica*, the most frequent *kdr* resistance mutations are S989 and L1014 (15). In *Ae. aegypti,* the 1014 codon requires at least two mutations to change to a *M. domestica* amino acid known to cause resistance; thus, the substitution L1014F, seen pervasively in *Anopheles* mosquitoes, has not been observed in this species (11). Instead, F1534C/L, V1016I/G, I1011V/M and V410L mutations have been associated with pyrethroid resistance in *Ae. aegypti* and confirmed experimentally (6). Other amino acid substitutions reported previously in *Ae. aegypti* include G923V, L982W, S989P, T1520I and D1763Y (11,16–18). Many of these mutations are often found in combination and appear only on specific continents. For example, V1016G and S989P appear limited to Asia, while V1016I has only been identified in the Americas and Africa and 723T only in the Americas (19).

The *ace-1* gene encodes acetylcholinesterase (AchE1), which is responsible for hydrolysis of acetylcholine terminating the transmission of neural signals. Organophosphates and carbamates bind to the acetylcholinesterase active site which inhibits hydrolysis and consequently neural signal termination, leading to insect death. Unlike mammals and some insects (including *Drosophila melanogaster*), mosquitoes usually have two copies of the *ace-1* gene. In *Anopheles* mosquitoes, the G119S amino acid substitution in *ace-1* is generally associated with resistance (all coordinates are based on *Torpedo californica*) (20,21). As with the *vgsc*, in *Ae. aegypti* such an amino acid change requires two mutations and has only been observed in one study in India (22). Despite the lack of described mutations in *ace-1*, resistance to organophosphates in *Aedes* is widespread in the Americas and Asia, while data from Africa is limited (6).

The *rdl* mutation is found in the γ-aminobutyric acid (GABA) receptor gene that controls neural signal inhibition through opening and closing of the transmembrane chloride channel on the cells of the mosquito nervous system. Cyclodienes (e.g., dieldrin) prevent interaction of GABA with its receptor, leading to neuron hyperexcitation and eventual insect death (23–26). The most common resistance mutation in this gene is A301S/G (*D. melanogaster* numbering) and is observed in multiple insects including mosquitoes of the *Anopheles* and *Aedes* genera (21,27). Despite a ban on the use of cyclodienes in 2001 (28) due to their slow degradation and environmental persistence, *rdl* mutations have persisted for decades later in vector populations, suggesting that they impart limited fitness costs (29,30).

Unlike *rdl*, *ace-1* and *vgsc,* which are targets of insecticides, the homodimer glutathione S-transferase (GST) is a detoxifying enzyme. Most organisms, including *Ae. aegypti*, have multiple GST enzymes of which epsilon two (GSTe2) has been associated with resistance to both DDT and pyrethroids (6,12,31,32). The *GSTe2* gene contributes to insecticide resistance through both enzyme overexpression and point mutations. Increased expression of this gene was linked to DDT resistance in *An. gambiae* (5,25,26,33) The L119F substitution in *GSTe2* was observed to enhance resistance to both DDT and pyrethroids in *An. funestus,* and I114T exacerbated resistance to DDT in *An. gambiae* (5,33–35). In *Ae. aegypti,* L111S and I150V mutations have been linked to temephos resistance *in silico* (36).

Despite observed phenotypic resistance of *Ae. aegypti* to all main insecticide classes across many countries in Africa, Americas, and Asia (6), the distribution of genetic variants in underlying candidate genes is less studied across *Aedes* populations compared to *Anopheles* species. Here, we examined a large (n=729), globally diverse dataset of publicly available *Ae. aegypti* whole genome sequencing (WGS) data to uncover the genetic diversity present in *vgsc, ace-1, rdl* and *GSTe2.* The diversity in insecticide resistance loci was interpreted alongside current global trends in phenotypic insecticide resistance in *Ae. aegypti*. This data provides a catalogue of genetic variants that could be involved in insecticide resistance and supports further studies on the molecular surveillance of emerging and spreading insecticide resistance mechanisms amongst *Ae. aegypti* populations.

## MATERIAL AND METHODS

### *Aedes aegypti* genomic data

We searched the NCBI SRA database for “*Aedes aegypti*” sample data and restricted results to WGS libraries where the number of bases contained implied at least 5-fold coverage when mapped to the reference genome AaegL5 (GCF_002204515.2) (32). We obtained a total of 703 WGS *Ae. aegypti* (non-AaegL5) libraries from 15 countries, across Africa (n=476, 8 countries), the Americas (n=191, 3 countries), Oceania (n=16, 1 country) and Asia (n=20, 1 country), and 26 colony samples of which 20 had known country of collection. Additionally, we included 7 *Ae. mascarensis* samples from Madagascar (n=4) and Mauritius (n=3) as outgroup (37–41) (**Table S1**).

### Insecticide resistance phenotypic data

Insecticide response data was only available for the Bora-Bora susceptible reference strain, which has been maintained in the insectary for 134 generations without any exposure to insecticides (42) and the Nakon Sawan reference strain, which is resistant to deltamethrin and temephos (41,43). Global insecticide resistance phenotype data was retrieved from the IR Mapper tool (44) (sourced on 19/04/2023), which covered 73 countries of which 8 overlap with samples in this study. No data was available for 5 countries (Kenya, Madagascar, Mauritius, South Africa, and Uganda); an additional literature search in PubMed failed to retrieve additional publicly available phenotypic data for *Ae. aegypti* in these countries. We included the data where the phenotype was tested with World Health Organization (WHO) tube or bottle bioassay or Centers for Disease Control and Prevention (CDC) bottle bioassay. Phenotypic data based solely on PCR or RT-PCR methods were excluded. Overall, we analysed 3,172 data points for 19 different insecticides across four insecticide classes (Pyrethroids, Organophosphates, Organochlorines and Carbamates) **(Table S2)**. Data points from IR mapper were reported as susceptible, possible resistance or resistant based on mortality as per WHO and CDC guidelines.

### Bioinformatic analysis

We aligned the WGS libraries using bowtie2 (v2.4.1) software (with a setting *--fast-local)* (45). We processed the alignment files using samtools (v1.7) software and SNPs were called using the GATK HaplotypeCaller tool (v4.1.9) with default settings (46,47). A minimum coverage of 5-fold was used to accept SNPs. We merged the individual VCF files into a multi-sample file using BCFtools (v1.9) (48). The impact of SNPs in the multi-sample VCF was predicted using snpEff software (v5.0) with AaegL5 genome annotation (GCF_002204515.2) (49). The alignment process was performed against the mRNA sequences of twenty *Ae. aegypti* genes **(Table 1)**. Four were loci linked to insecticide resistance [*vgsc* (XM_021852340.1), *rdl* (XM_021840622.1), *ace-1* (XM_021851332.1) and *GSTe2* (XM_021846286.1)] and the remaining sixteen genes were used to establish population structure. One of these was mitochondrial *cox1* (YP_009389261.1) and the remaining fifteen genes were evenly spread across all three *Ae. aegypti* chromosomes **(Table 1)**. These 15 genes were determined to have unique genome-wide exon sequences (using NCBI BLASTn v2.9.0 with *--word-size* 28 and *--evalue* 0.01) which minimised potential mis-mapping of WGS reads to the *Ae. aegypti* genome known to contain many duplications (50). Read coverage per nucleotide per gene was calculated using the samtools “depth” function and was used to identify possible gene duplications in samples (48). We merged the coverage data into a single data matrix and removed all regions except gene exons, because intronic regions contained high numbers of repeats. For each sample, we divided each per base coverage value by that sample’s overall median coverage across all genes, except *vgsc* and *GSTe2,* which may have copy number variants. We applied UMAP (v0.5.1) software (with a *Euclidean* distance metric) on this scaled matrix to identify gene clusters based purely on the coverage (51).

**Table 1.** The genes analysed. * - The annotation in GCF_002204515.2 assembly has missing start codon for mitochondrial *cox1* and as a result snpEff did not distinguish between synonymous and non-synonymous SNPs. CDS = coding sequence

| Gene | Product | Chr | CDS Len | Resistance gene | Unique missense SNPs | Unique synonymous SNP |
| --- | --- | --- | --- | --- | --- | --- |
| XM_021851049.1 | TATAmodulator | NC_035107.1 | 3178 |  | 557 | 1040 |
| XM_001648700.2 | YdcI | NC_035107.1 | 2902 |  | 380 | 602 |
| XM_021851750.1 | LOC110678629 | NC_035107.1 | 1095 |  | 35 | 13 |
| XM_021857384.1 | LOC5580295 | NC_035107.1 | 2479 |  | 657 | 437 |
| XM_001652683.2 | PotassiumChannel | NC_035107.1 | 1017 |  | 39 | 262 |
| XM_021840622.1 | GABA | NC_035108.1 | 1653 | Yes | 64 | 180 |
| XM_021841341.1 | AngiogenicFactor | NC_035108.1 | 1811 |  | 334 | 439 |
| XM_001664194.2 | TIFIID2 | NC_035108.1 | 3755 |  | 217 | 738 |
| XM_001662595.2 | Mcm6 | NC_035108.1 | 2429 |  | 99 | 167 |
| XM_001657120.2 | Cytochrome b-c1 | NC_035108.1 | 797 |  | 48 | 122 |
| XM_021846286.1 | GSTE2 | NC_035108.1 | 666 | Yes | 109 | 158 |
| XM_021847043.1 | Carbohydratesulfotransferase | NC_035108.1 | 1330 |  | 194 | 291 |
| XM_001657462.3 | LOC5567548 | NC_035109.1 | 1745 |  | 334 | 396 |
| XM_021850261.1 | ZincFinger | NC_035109.1 | 1497 |  | 235 | 267 |
| XM_021851332.1 | ACE1 | NC_035109.1 | 2102 | Yes | 99 | 144 |
| XM_001649087.2 | grpE | NC_035109.1 | 676 |  | 63 | 59 |
| XM_001649790.2 | LOC5565494 | NC_035109.1 | 2445 |  | 527 | 535 |
| XM_021852340.1 | VGSC | NC_035109.1 | 6379 | Yes | 202 | 873 |
| XM_021853012.1 | LOC5579101 | NC_035109.1 | 1836 |  | 378 | 208 |
| YP_009389261.1 | COX1 | NC_035159.1 | 1536 |  | 1230* | 0* |

### Population genetics analysis

To determine population structure, we used UMAP software (with *Russell-Rao* distance metric) on the multi-sample VCF, followed by application of HDBSCAN (v0.8.28) (51,52) to determine sample clustering (see (53–55) for recent applications). This work was performed in python (v3.7.6), with scripts available from https://github.com/AntonS-bio/resistance-AedesAegypti. Linkage disequilibrium was calculated using vcftools on phased vcf files created with beagle (v 22Jul22.46e) software to provide a R^2^ value for each combination of non-synonymous mutations by sample country. Plots of these values were visualised using the gaston (v1.5.9) package in R.

### Protein structure modelling

Protein structure modelling was performed using AlphaFold Multimer software with full protein databases (56,57). When referring to substitutions and their effects on proteins, we have followed the established nomenclature based on reference resistance linked proteins and structures in the protein databank: ACE1 (2C4H; *Tetronacre californica*), GABA receptor (NP_729462.2; *Drosophila melanogaster*), GSTe2 (XP_319968.3; *An. gambiae*) and VGSC (NP_001273814.1; *Musca domestica*) (58,59). Unless otherwise specified, all substitution coordinates are with respect to these reference sequences.

## RESULTS

### Genetic variation and population structure

Across the 729 *Aedes* samples from 15 countries, a total of 1,829 SNPs (474 non-synonymous (NS)) were detected across the CDS of four insecticide resistance associated genes (*vgsc*, *rdl*, *ace-1* and *GSTe2*), and 9,673 SNPs were identified across the CDS of 15 non-resistance associated genome-wide gene (**Table 1**, **Table 2, Table S3**).

**Table 2.**
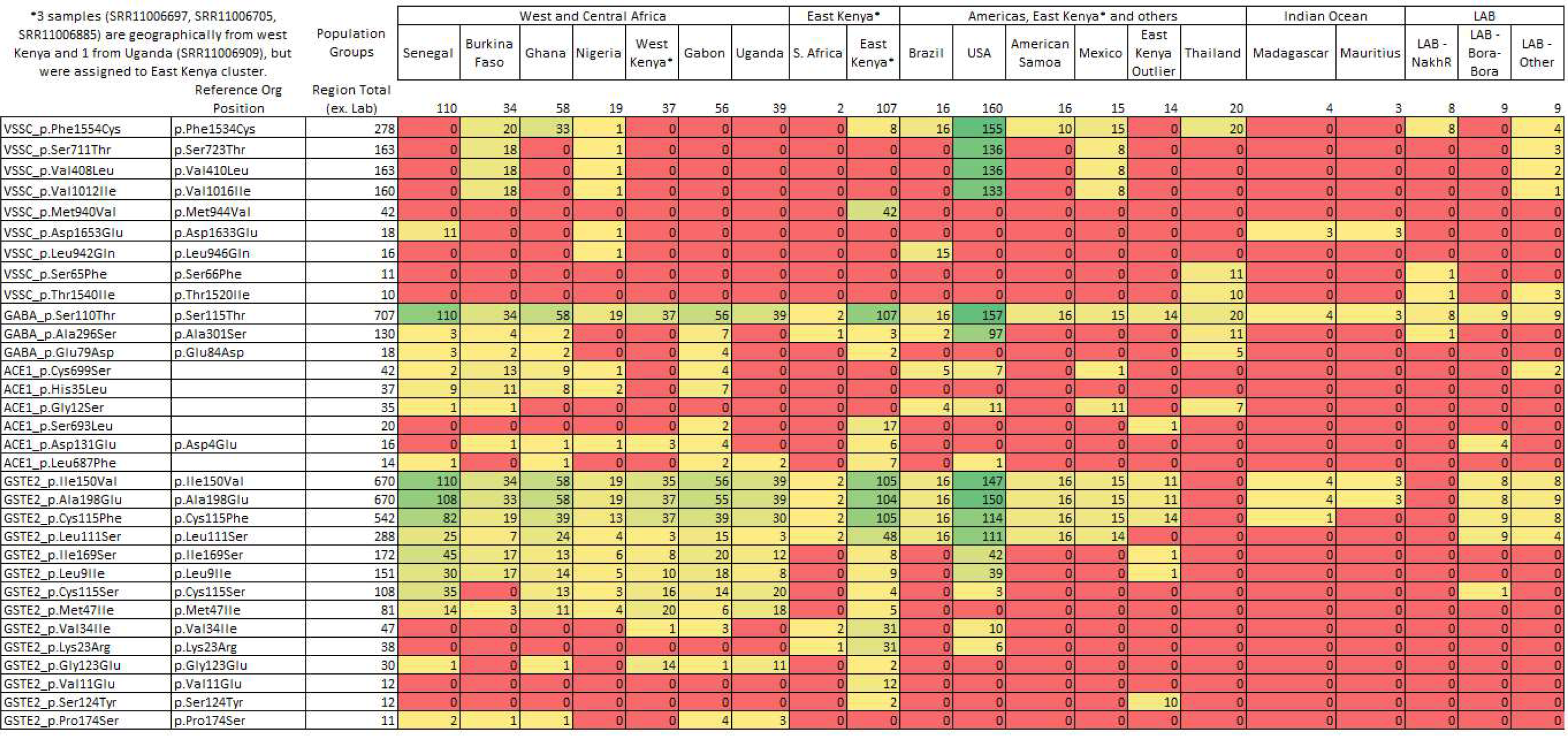
Missense mutations identified in samples and occurring in more than 10 non-lab sample. The full list of mutations is available in Supplementary Table B.

Using the SNPs from the CDS of 15 genes not associated with insecticide resistance, a UMAP clustering analysis revealed five distinct clusters **(Figure 1(A))**, broadly linked to: (i) eastern Kenya and South Africa (n=112); (ii) west, central Africa and west Kenya (n=350); (iii) the Americas, Thailand, and other (n=258); (d) the Bora-Bora mosquito line from French Polynesia (n=9); (e) *Ae. mascarensis* from Madagascar and Mauritius (n=7). Similar results were obtained when analysing only the 1,829 SNPs in genes that are associated with resistance (**Figure 1(B)**). These results are broadly consistent with previous reported population structure of *Ae. aegypti* using SNPs and microsatellite data, where African samples formed one cluster and samples from Asia, America and the Caribbean comprised another cluster (60). As we observed a separation of most eastern Kenyan samples (n=121) from west Kenya (n=37), we investigated the genotype data in these groups independently. Some eastern Kenyan samples (n=14/121) from a human-biting colony of domestic *Ae. aegypti,* originally collected indoors in Rabai (60,61), clustered with non-African samples (Americas and Thailand and other cluster), as previously observed. When including only non-African samples, the UMAP clustering analysis revealed modest separation of the samples from Brazil, Mexico, French Polynesia, American Samoa and Thailand. For the samples from Africa, clustering separated east Kenyan samples from the rest **(Figure S1)**. The same patterns were detected across both resistance and non-resistance genes (**Figure S1**). Clustering using mitochondrial *cox1* gene was different from the results based on chromosomal loci (**Figure 1(C-F))**. In multiple samples, SNPs had heterozygous *cox1* genotypes possibly multiploidy due to the presence of previously described copies of nuclear mitochondrial (NUMT) DNA which could confound clustering (62,63).

**Figure 1.**
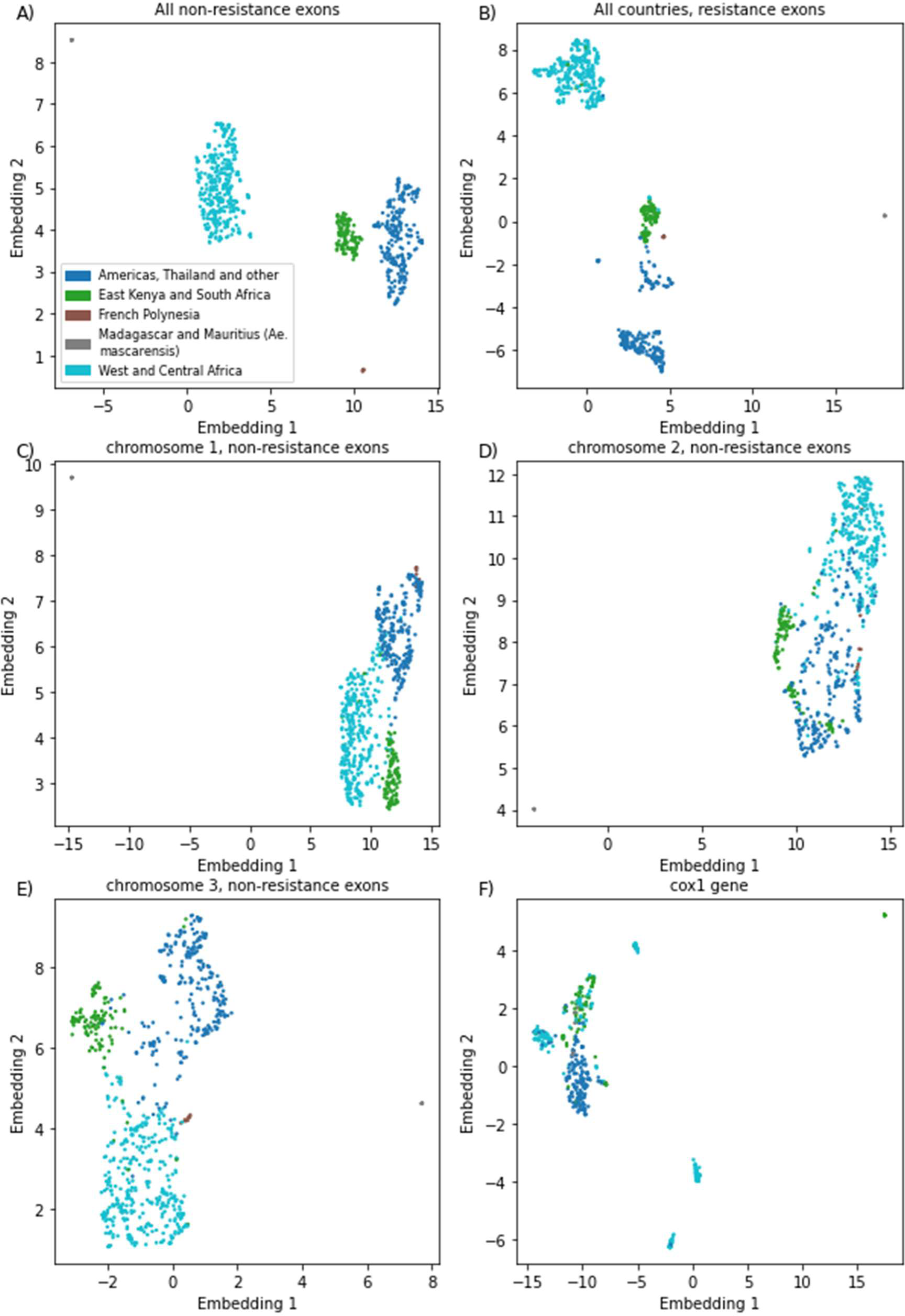
Population structure using UMAP embedding of SNPs from non-resistance linked genes (A), (C), (D), (E), and resistance linked genes (B) and *cox1* (F).

### Genetic variation across insecticide resistance associated genes

#### Vgsc

In the *vgsc* gene, a total of 1075 SNPs (202 non-synonymous; NS) were identified, of which 36 NS SNPs were present in >1 sample, including eight mutations previously linked to insecticide resistance (V410L, G923V, S989P, I1011M, V1016I/G, T1520I and F1534C) **(Table 2, Table S3)**. We did not observe any other pyrethroid resistance associated substitutions such as L982W, detected previously in Vietnam and Cambodia, and D1763Y reported in Taiwan. However, the D1763G mutation was present in a single USA sample (11,16–18). The most frequent mutations were F1534C (39%), S723T (23%), V410L (22%) and V1016I (22%) (**Figure 2**). The most prevalent F1534C mutations occurred in nearly all samples from the Americas (186/191) and Thailand (20/20). The frequency of F1534C was lower in African samples, appearing only in Burkina Faso (n=20/34), Ghana (n=33/58), Nigeria (n=1/19) and East Kenya (n=8/107). The F1534C mutation was accompanied by V1016I, S723T and V410L substitutions in most samples from USA, Burkina Faso, and Mexico, as well as in a single Nigerian sample. In Thailand, F1534C co-occurred in many samples with V1016G, T1520I and S989P **(Table 2)**.

**Figure 2.**
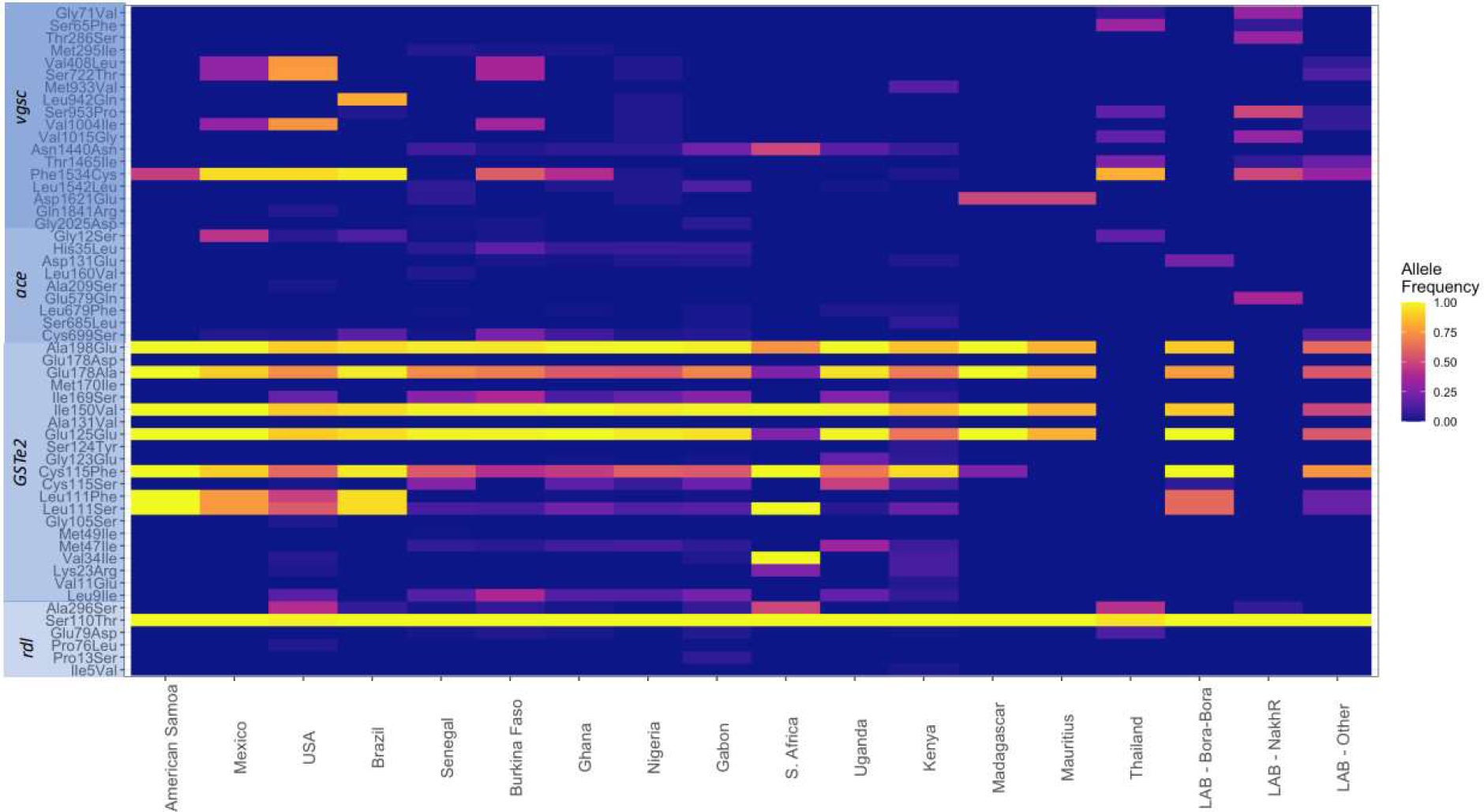
Allele frequency of each missense SNP across the insecticide resistance associated genes; *vgsc, ace-1, rdl, and GSTe2*, by country. Only SNPs with at least 10 samples with a non-reference allele are shown. mutation.

Several mutations were found to be regionally specific. The V1016G mutation was found only in Asia (Thailand) while V1016I was detected in USA, Mexico, and a few countries in Africa (19)). The M944V substitution was unique to East Kenya (n=42/107), L946G was almost exclusive to Brazil (n=15/16) except for one Nigerian sample. The V1016G, T1520I (n=10/20), S989P (n=7/20), and S66F (n=11/20) were also almost exclusive to Thailand, apart from a single Nigerian and a Brazilian sample **(Table 2)**. Two conservative in-frame insertions occurred in ∼20% of west and central African samples, which included an addition of amino acid Glycine (Gly) into a sequence of four consecutive Gly (positions 2047-2050), and an addition of Serine-Glycine (positions 2016 and 2017).

#### *Rdl* (GABA receptor)

In the *rdl* gene, we identified a total of 244 SNPs (64 NS), of which only 17 NS SNPs occurred in >1 sample and the most frequent were G84A, S115T and A301S. The S115T substitution was present in almost all samples (n=733/736) including all *Ae. mascarensis*. (**Figure 2**, **Table 2**). The T115 is the dominant allele in *An. gambiae* suggesting that the common ancestor of both *An. gambiae* and *Ae. aegypti* had the 115T allele, and a mutation in the *Ae. aegypti* reference strain changed T to S (64).

The previously described A301S substitution, associated with resistance to organochlorines, was frequent in the USA (n=97/160) and Thailand (n=11/20), and infrequent in a few countries in Africa (**Table 2**) (21,27). This substitution is located on the a-helix forming the protein pore **(Figure S2)**. The only other notable mutation was E84D present in 18 samples (Africa n=13, Thailand n=5), and located on the outward facing section of the protein but could not be robustly modelled by the AlphaFold software.

#### Ace-1

A total of 243 SNPs were identified in the *ace-1* gene, of which 99 led to amino-acid substitutions, with 30 present in >1 sample **(Table 2)**. Only 6 amino-acid substitutions (G12S, H35L, D131Q, L687F, S693A, C699S) occurred in >10 samples (**Figure 2**). The most frequent mutation was C699S (n=42/736), which was present in samples from west and central Africa (n=29) and the Americas (n=13). The second most frequent substitution was H35L (5.0%) observed only in west and central African samples. The third most frequent substitution was G12S (4.8%) found mostly in the Americas (n=26/37) and Thailand (n=7/37) (**Table 2**). All three substitutions are defined in *Ae. Aegypti* coordinates because these amino acids are outside the range of the *T. californica* reference ACE1 (PDB: 2C4H). In fact, only 20 substitutions had a corresponding coordinate in the *T. californica* protein **(Table 2**). The only substitution in *Ae. mascarensis* was T55P (*T. californica* coordinates) present in all samples of this species. We modelled the ACE1 protein structure in AlphaFold, and in line with results of crystallographic experiments, the residues 1-131 and 660-702 were disordered, likely reflecting their role in anchoring the protein to the cellular membrane and receptor proteins (65). The G119S resistance substitution commonly reported in ACE1 in other insect species was not detected in this dataset. This absence is likely because G119S would require two nucleotide substitutions in *Ae. aegypti*. Further, instead of two *ace* genes commonly found in insects, the *Ae. aegypti* reference genome has four *ace* genes including one analysed here (LOC5578456) and three others (LOC5574466, LOC5575867, LOC5570776). The mRNA encoding the cognate proteins had <5% pair-wise coverage which rules out recent duplication as the origin of these genes. One of these loci (LOC5570776) had the 119S amino acid. We found that despite the very high prevalence of transposable elements in *Ae. aegypti,* this gene remains uninterrupted by them suggesting this locus might be functional (32).

### GSTe2

The *GSTe2* gene has a variable copy number in *Ae. aegypti,* and the reference genome contains four copies of this gene (32). The variable copy number was also evident in our analysis. Because we used short read data, we could not robustly assign each mutation to individual *GSTe2* loci. A total of 267 SNPs were detected in *GSTe2* genes, with 109 leading to amino-acid substitutions, of which 42 were present in >1 sample **(Table 2)**. Seven substitutions were highly frequent: I150V (n=670), A198E (n=670), C115F (n=542), L111S (n=288), I169S (n=172), L9I (n=151) and C115S (n=108) **(Figure 2)**. The samples from Thailand had neither synonymous nor missense mutations in *GSTe2,* which we confirmed by visual examination of the read alignments. The C115F substitution was present in almost all countries (except Thailand and Mauritius). The C115S substitution was most common in Africa (n=101/353). In addition to C115F/S, we observed two other common substitutions (L111S, L9I) at the DDT binding site (66). The L111S substitution (n=288/736) appears globally distributed, and L9I was found mainly in Africa and USA, but not observed in *Ae. mascarensis*. The I169S mutation was common in the presence of L9I. Based on a high confidence AlphaFold protein structure model for GSTe2, the I169S mutation is not part of either glutathione or DDT binding site; however, it interacts with both F115 and M111, which are part of the glutathione binding pocket (**Figure S3**).

### Gene duplications

Gene variable copy numbers were identified based on excess median-scaled read coverage. For the *vgsc* gene, a group of 26 samples had potential duplications, with a median-scaled coverage of 1.4-fold compared to 1.0-fold for the rest of the samples. The samples in this set came from a disparate group of countries: Senegal (n=13), American Samoa (n=4), and USA (n=3), Mexico (n=2), Mauritius (n=2), Kenya (n=1) and Thailand (n=1) **(Table S1)**.

For *GSTe2*, two groups of samples had likely copy number events. First, a group of samples with median 4.2-fold median-scaled coverage consisting of samples from Thailand (n=27/28) including samples from the Nakh lab strain, USA (n=38/160), Mexico (n=5/15), Brazil (n=1/16) and two from the Vienna F4 colony (67). A second group consisted of samples from USA (n=15/160) and Mexico (n=9/16) with median-scaled coverage of 9.3-fold compared to 0.9-fold for the rest of the samples **(Table S1, Figure S4).** In our search of the literature, we did not identity previous reports of such high duplication rate; this finding requires further validation. However, this result also shows that majority of *Ae. aegypti* reference sequence have single copy of *GSTe2*, in contrast to the reference strain which has four (32).

### Linkage disequilibrium between missense mutations

We examined the geographical distribution of the non-synonymous SNPs across the four resistance genes and observed that many mutations co-occur together in certain populations (**Figure 2**). For each locus, per population, we assessed the pairwise linkage disequilibrium (LD) of non-synonimous SNPs. We found twenty-seven pairwise SNPs that had, without adjusting for multiple testing, an R^2^ value above 0.5 (*GSTe2 n=15*, *vgsc n=9*, *ace-1 n=2,* and *rdl n=1*) **(Table S4**). The *GSTe2* mutations L9I/I169S (Burkina Faso, Kenya, Gabon, Ghana, Uganda) and I150V/A198E (Kenya, French Polynesia, Mauritius) were detected with a R^2^ >0.5 in several countries. In the *vgsc* gene, several SNPs that have been associated with insecticide resistance also had R^2^ > 0.5, particularly V410L, V1016I, V1016G and F1534C.

### Geographical distribution of Insecticide Resistance Mutations and Phenotypes

The IR mapper was used to obtain phenotypic data for 8 of the 15 countries examined in this study. These phenotypes show disparity between the availability of phenotypic and genomic data, for example, Brazil and Thailand have the highest number of bioassay records while only having 16 and 20 genomic sequences available, respectively. However, in some countries there was genomic data available with limited phenotypic data, such as Uganda and Kenya. Phenotypic data available for each country from IR Mapper was mapped to the co-occurrence of nine mutations previously associated with insecticide resistance (A301S (RDL) associated with organochlorine resistance, and F1534C, T1520I, V1016I/G, I1011V/M, S989P, G923V, V410L (VGSC) all associated with pyrethroid resistance). Thailand, Burkina Faso, and the USA had the highest proportion of samples with several known insecticide resistance mutations **(Figure 3)**. This is supported by the Thailand phenotypic data from IR Mapper, which shows reports of resistance to all four main insecticide classes in this country (Figure 4), particularly to organochlorines, carbamates and pyrethroids. Elevated levels of resistance have also been reported in southeast Asian regions, such as Indonesia, Malaysia, and Thailand; however, there are gaps in the genomic data from these countries (68–71). For the USA there is no information on phenotype data on IR Mapper, but resistance to pyrethroids has been reported in several states (72–74).

**Figure 3.**
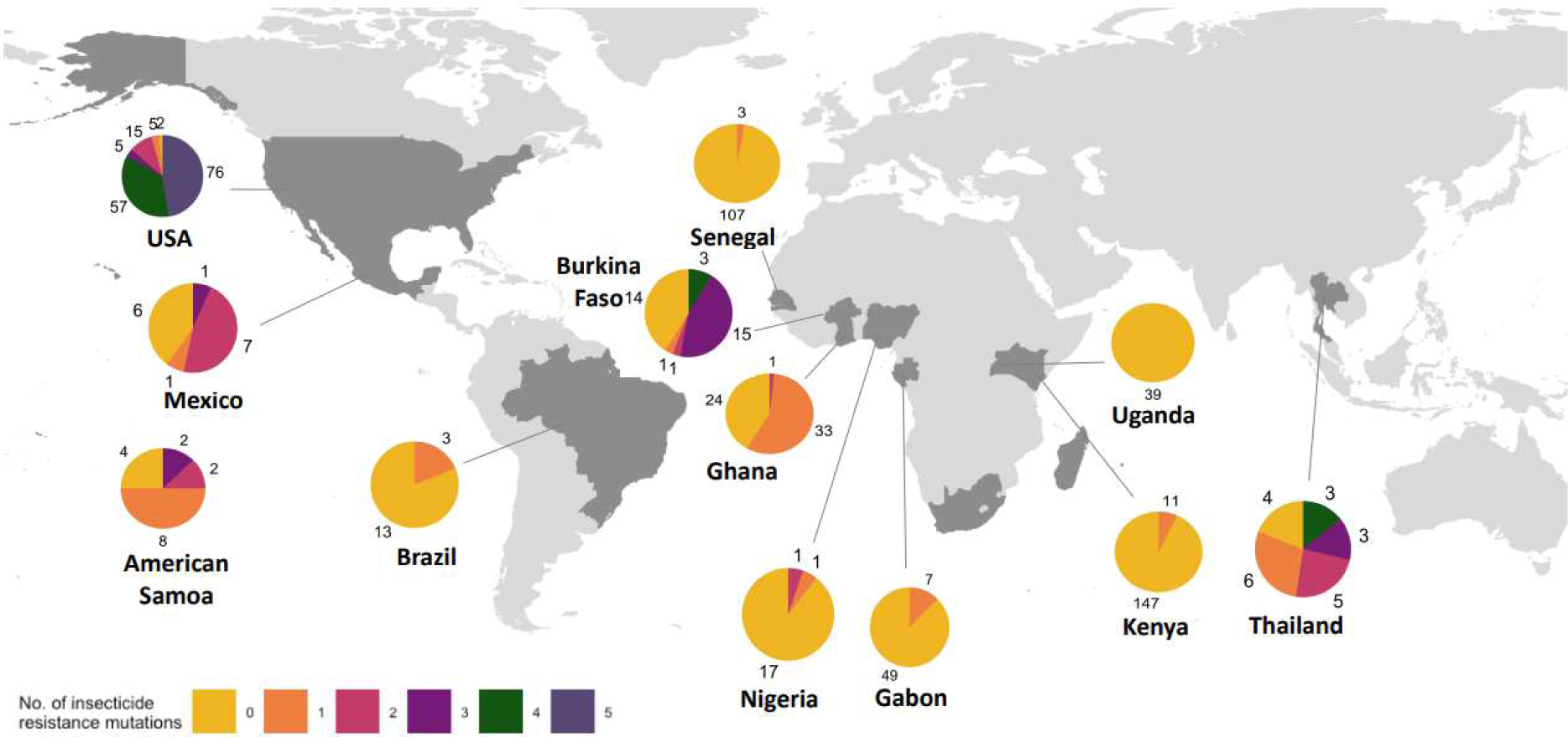
Proportion of samples with 1 or more mutations associated with insecticide resistance in each geographical population. Insecticide resistance SNPs included are: A301S *(rdl*), F1534L/C, T1520I, V1016I/G, I1011V/M, S989P, G923V, V410L (v*gsc*). Only populations with more than 10 samples were included.

**Figure 4.**
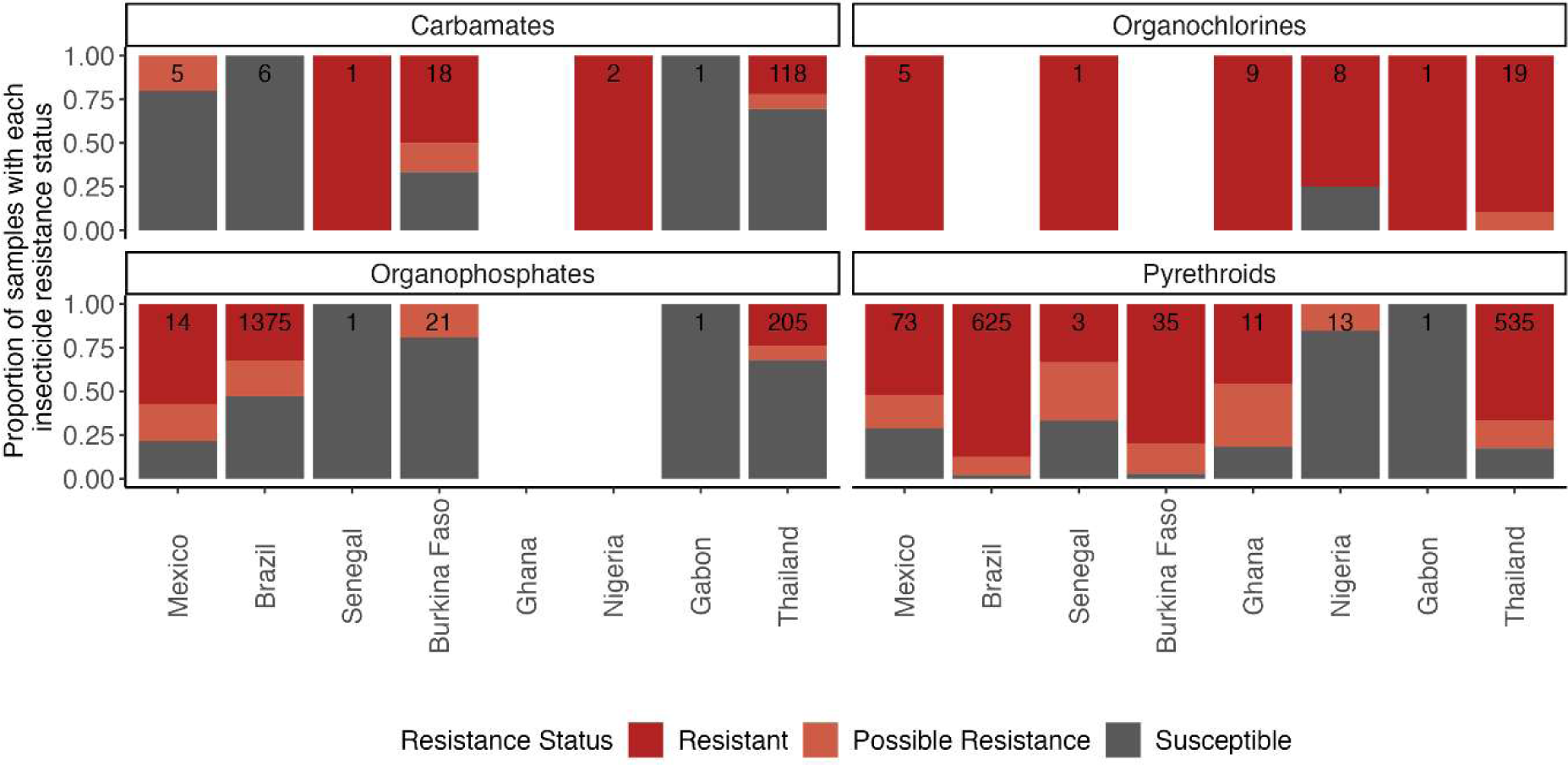
Publicly available phenotype data for *Ae. aegypti* showing the proportion of records that report resistance, possible resistance and susceptibility. Numbers denote total number of records for the insecticide class for that country region (44). Only data collected on *Aedes aegypti* after 2000 were included for countries that were present in the WGS data set.

In Africa, 53% of samples from Burkina Faso had more than two insecticide resistance mutations, all in the *vgsc* gene. Burkina Faso also had the highest reported resistance to pyrethroids when compared to the other African samples in this data set (Nigeria, Senegal, Ghana, and Gabon). Levels of resistance to pyrethroids varied between the 8 countries analysed here. The highest levels of resistance were also observed in Brazil, Mexico, and Thailand, coinciding with samples with the most mutations in the *vgsc* gene (excluding the USA, where limited phenotypic data is available) (**Figure 3**, **Figure 4**).

The data from IR mapper showed that the largest number of reports of resistance involved insecticides of the organochlorine class. Mutations associated with this resistance include SNPs in the *vgsc* and *rdl* genes. However, countries with high resistance to organochlorines, such as Senegal and Nigeria have no or very low frequency of mutations in these loci. As the genomic data presented here do not have matching phenotypic information, it is possible that these samples were from a susceptible background or that there are other mechanism of resistance causing the observed phenotype. The least resistance was reported against organophosphates, although resistance is still high in Mexico, followed by Brazil and Thailand **(Table 2)**. These countries only have 1 mutation, G12S, in the *ace* gene common across all of them.

## DISCUSSION

We explored the genetic diversity present in four genes (*vgsc, ace-1, rdl* and *GSTe2)* involved in insecticide response across 729 *Ae. aegypti* and 7 *Ae. mascarensis* samples from 15 countries. We identified many known and unreported amino-acid substitutions which may be involved in insecticide resistance. This catalogue of genetic variants is a valuable resource that can be explored to investigate molecular mechanism associated with insecticide resistance together with phenotypic information and used to design diagnostics genetic markers for molecular surveillance.

The populations with greater numbers of amino acid substitutions linked to insecticide resistance were Thailand (RDL: A301S; VGSC: V410L, S989P, V1016G and F1534C) and the USA (RDL A301S; VGSC: V410L, Gly923V, I1011M and F1534C). In Africa, the substitutions most frequently observed were RDL A301S and VGSC V410L and F1534C, but many countries had none of the reported mutations. We have also observed that VGSC V410L and S723T co-occur in all but one sample. None of the Thai samples had any mutations in the *GSTe2* gene, despite having adequate read coverage. In other countries, we detected two common mutations in GSTe2 (C115F/S and L111S/F) in the DDT binding site. The C115F and C115S mutations were most frequent in Kenya (n=142, n= 20), the USA (n=114, n = 20) and Senegal (n=82, n = 35). Previous work involving DDT docking with *An. gambiae* GSTe2 has suggested that one of the DDT’s planar p-chlorophenyl rings can fit into a sub-pocket, but the other ring faces spatial hindrance from M111 and F115 in the side chains (66). In *An. gambiae*, the M111S substitution would require two nucleotide changes in contrast to one required for L111S/F in *Ae. aegypti*. To our knowledge, there are no reports of *An. gambiae* M111S or F115C/S; although the latter substitution requires a single amino acid change. These two substitutions were detected in almost all countries in this *Aedes* dataset.

We found only two mutations on the surface of the ACE1 pocket directly involved in hydrolysis (A81S, n=5; D85H, n=2) (13). Since we did not have phenotype data, we cannot determine if these mutations are associated with resistance, but their low prevalence would appear at odds with much higher rate and multiple instances of emergence of G119S in *An. gambiae* (75). Nevertheless, further functional work can contribute to elucidating the involvement of these mutations in resistance phenotypes.

We have also explored the possibility of gene duplications, and detected such variants in *GSTe2* in USA, Mexico, Brazil, and Thailand, which are of interest due to the high rates of permethrin resistance reported in the Americas and Asia (76,77). We found no duplications in west and central Africa or Eastern Kenya and South Africa regions (6), but bioassay data in these regions is lacking. The possible duplication of the gene encoding VGSC is more puzzling. Previous research in *D. melanogaster* found that individuals lacking VGSC are not viable, but in contrast those with a single functioning gene copy are healthy apart from increased temperature sensitivity (78). However, DDT and pyrethroids both prolong the open state of VGSC, so the extra gene copy is unlikely to induce resistance through increased number of pores (14). Experimental work is required to explain the functional role of the extra copy and determine if it is associated with increased insecticide resistance. Long-read sequencing can help to validate the duplications detected and the differences between the *vgsc* sequences.

The inferred population structure was broadly consistent with previous research based on chromosomal loci. We even identified the two previously described distinct subpopulations of *Ae. aegypti* in Rabai District of Kenya (60). An important observation for future research is that the *cox1* gene and other mitochondrial loci may be problematic for population studies in *Ae. aegypti* because of the unknown number of *cox1* copies per genome (62,63). This is the result of unknown numbers of mitochondria per cell, unknown number of mitochondrial DNA copies on chromosomes, and unknown allelic diversity of all these *cox1* sequences.

While we focused on exploring the genetic diversity in four genes associated with target site insecticide resistance, there are many loci that could have an important role, particularly in metabolic resistance. Multiple P450 genes, particularly members of the CYP6 and CYP9 subfamilies, have been associated with resistance by overexpression when comparing insecticide-resistant to susceptible strains (79–81).

Having both phenotypic and genotypic data is fundamental for the full understanding of the link between phenotypic resistance and genetic mutations, as well as cross resistance mechanisms. Unfortunately, we did not have phenotypic data for all the countries with genotypic data in this study. We strongly advocate that where possible, phenotypic data be generated for samples with genomic sequences.

Further work on exploring genetic diversity in these gene families, particularly using long-read sequencing to support assembly and correct assignment of copy numbers to each individual gene, may reveal important molecular markers that can be involved in insecticide resistance. Genomic studies, like ours, can provide guidance to functional studies and inform the design of genotyping assays for large scale surveillance of insecticide resistance.

## Supporting information

Oversized supplementary tables

## ACKNOWLEDGEMENTS

E.L.C. is funded by an MRC LID PhD studentship TGC and SC are funded by UKRI grants (BBSRC BB/X018156/1; MRC MR/X005895/1; EPSRC EP/Y018842/1)).

## AUTHOR CONTRIBUTIONS

AS, SC and TC designed the study. AS and EC analysed the data under the supervision of TC and SC. All authors interpreted the results. AS and EC wrote the first draft of the manuscript. All authors have edited and approved the final version of the manuscript.

## SUPPLEMENTARY FIGURES AND TABLES

**Supplementary Figure 1.**
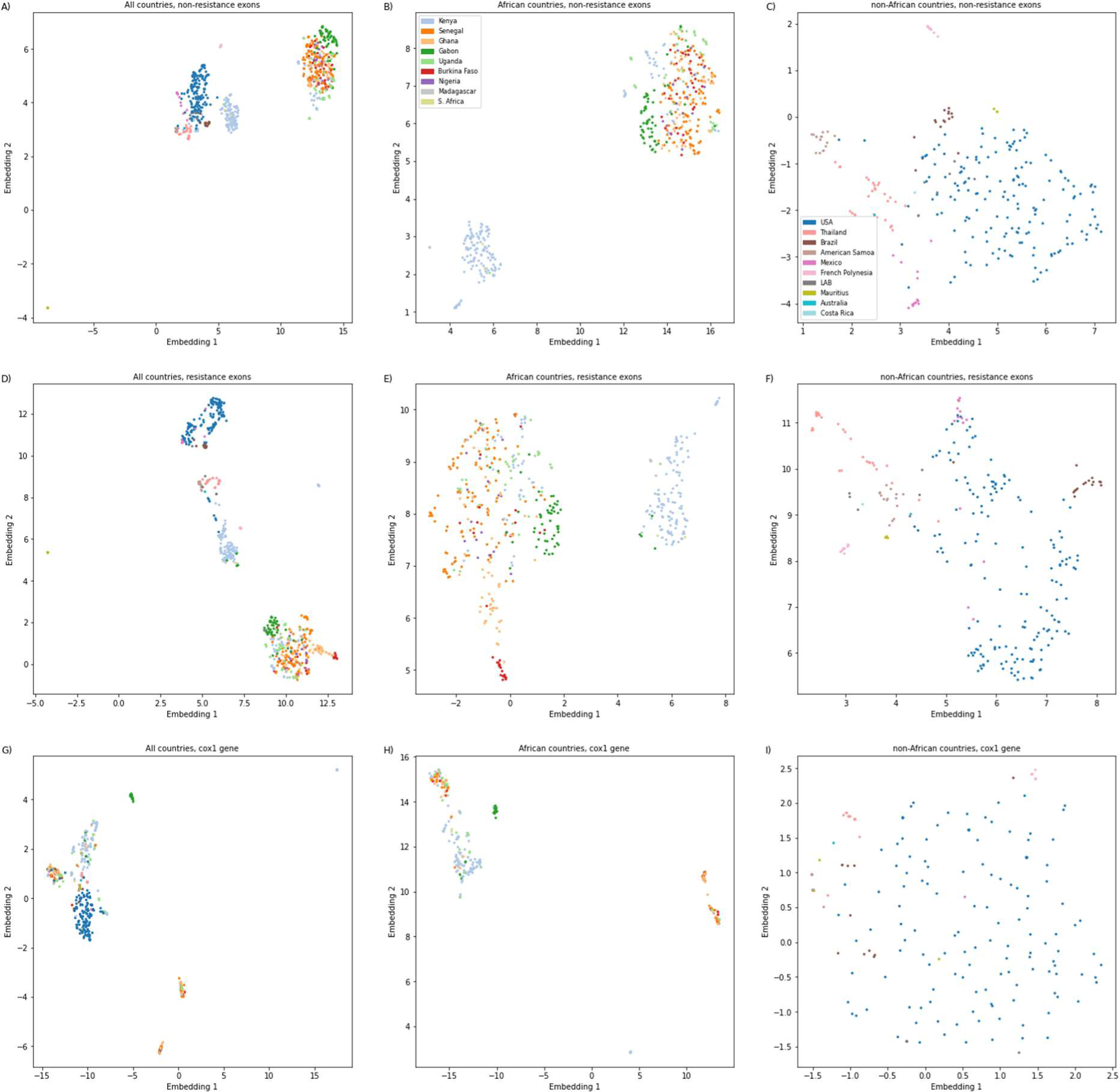
Population structure using UMAP embedding of SNPs for different geographical regions.

**Supplementary Figure 2.**
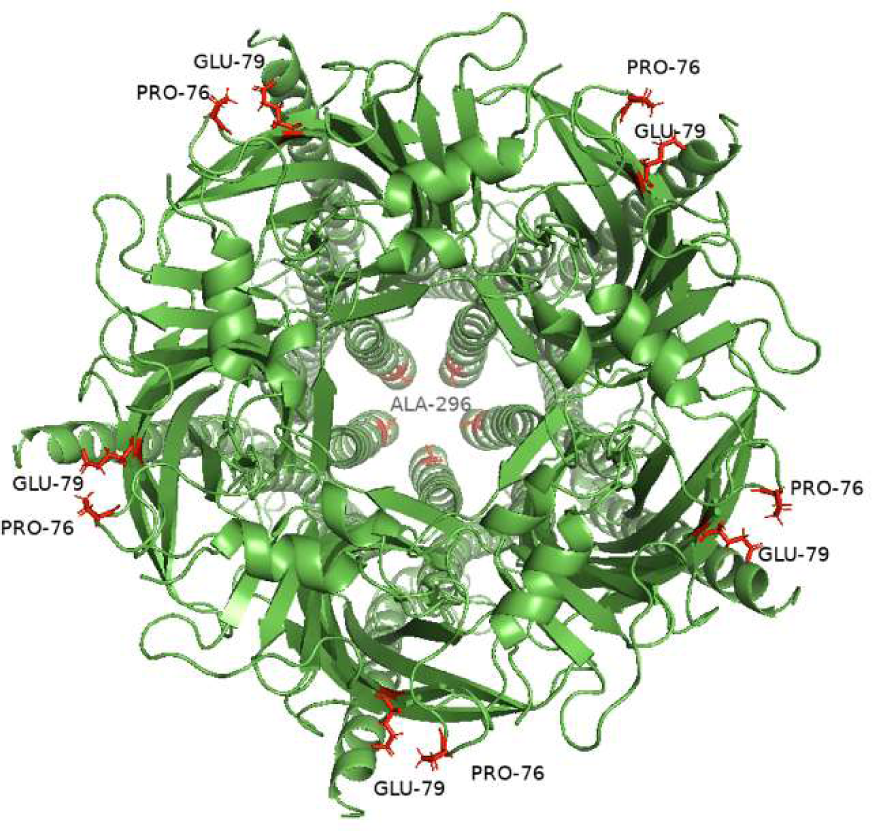
GABA receptor protein structure including mutations found in >10 isolates.

**Supplementary Figure 3.**
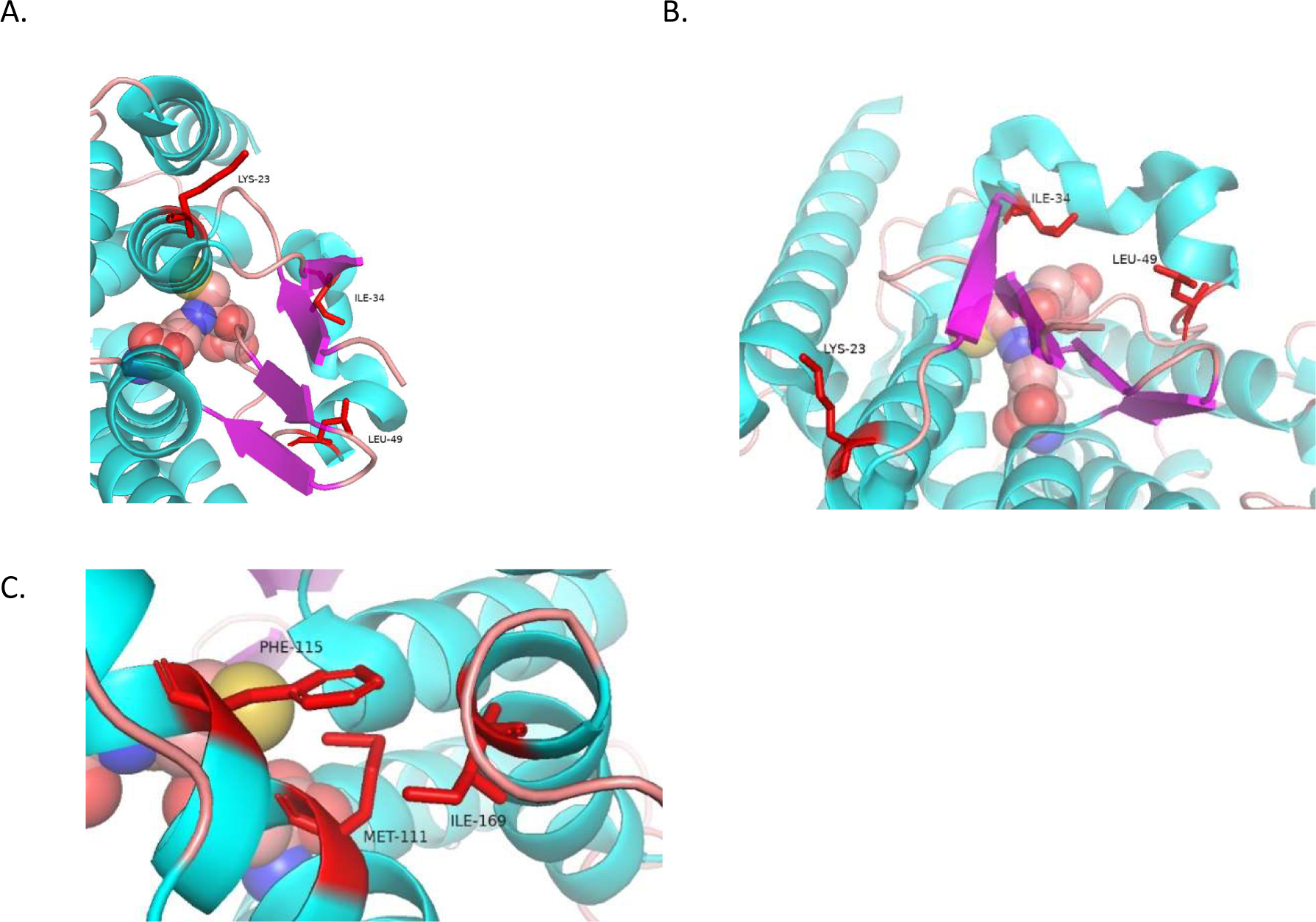
*GSTe2* mutations specific to East Kenya and South Africa (A,B) and common substitutions (Cys115Phe/Ser and Leu111Ser) together with west and central Africa specific Ile169Ser substitution. The residue at position 111 is methionine because we used PDB 2IMI structure of *An. gambiae* to show accurate ligand docking (**66**).

**Supplementary Figure 4.**
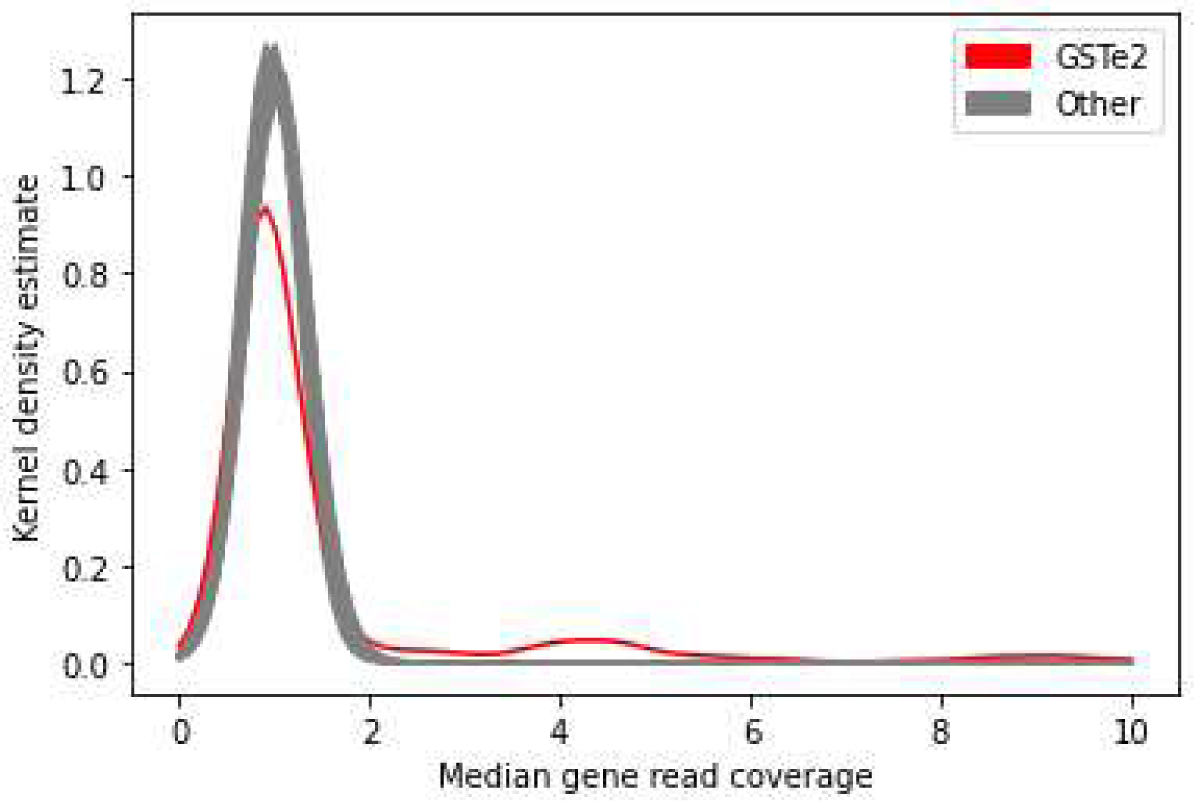
Median per-base read coverage across samples for GSTe2 and other genes. The coverage was normalised for each sample using median coverage across the genes for that sample. Two peaks are visible in GSTe2 at 4 and 9 median gene read coverage.

## REFERENCES

1. Lwande OW, Obanda V, Lindström A, Ahlm C, Evander M, Näslund J, et al. Globe-Trotting Aedes aegypti and Aedes albopictus: Risk Factors for Arbovirus Pandemics. Vol. 20, Vector-Borne and Zoonotic Diseases. Mary Ann Liebert Inc.; 2020. p. 71–81.

2. Kraemer MUG, Sinka ME, Duda KA, Mylne AQN, Shearer FM, Barker CM, et al. The global distribution of the arbovirus vectors Aedes aegypti and Ae. Albopictus. Elife. 2015 Jun 30;4(JUNE2015).

3. Brown JE, Evans BR, Zheng W, Obas V, Barrera-Martinez L, Egizi A, et al. Human impacts have shaped historical and recent evolution in Aedes aegypti, the dengue and yellow fever mosquito. Evolution. 2014 Feb;68(2):514–25.

4. Rocklöv J, Dubrow R. Climate change: an enduring challenge for vector-borne disease prevention and control. Nat Immunol. 2020 May 1;21(5):479–83.

5. Liu N. Insecticide Resistance in Mosquitoes: Impact, Mechanisms, and Research Directions. 101146/annurev-ento-010814-020828. 2015 Jan 71;60:537–59.

6. Moyes CL, Vontas J, Martins AJ, Ng LC, Koou SY, Dusfour I, et al. Contemporary status of insecticide resistance in the major Aedes vectors of arboviruses infecting humans. PLoS Negl Trop Dis. 2017 Jul 20;11(7):e0005625.

7. Ingham VA, Wagstaff S, Ranson H. Transcriptomic meta-signatures identified in Anopheles gambiae populations reveal previously undetected insecticide resistance mechanisms. Nat Commun. 2018 Dec 1;9(1).

8. Ingham VA, Anthousi A, Douris V, Harding NJ, Lycett G, Morris M, et al. A sensory appendage protein protects malaria vectors from pyrethroids. Nature. 2020 Jan 16;577(7790):376–80.

9. Messenger LA, Impoinvil LM, Derilus D, Yewhalaw D, Irish S, Lenhart A. A whole transcriptomic approach provides novel insights into the molecular basis of organophosphate and pyrethroid resistance in Anopheles arabiensis from Ethiopia. Insect Biochem Mol Biol. 2021 Dec 1;139.

10. Balabanidou V, Kampouraki A, Maclean M, Blomquist GJ, Tittiger C, Juárez MP, et al. Cytochrome P450 associated with insecticide resistance catalyzes cuticular hydrocarbon production in Anopheles gambiae. Proc Natl Acad Sci U S A. 2016 Aug 16;113(33):9268–73.

11. Du Y, Nomura Y, Zhorov BS, Dong K. Sodium Channel Mutations and Pyrethroid Resistance in Aedes aegypti. Insects. 2016 Dec 1;7(4).

12. Pavlidi N, Vontas J, Van Leeuwen T. The role of glutathione S-transferases (GSTs) in insecticide resistance in crop pests and disease vectors. Curr Opin Insect Sci. 2018 Jun 1;27:97–102.

13. Cheung J, Mahmood A, Kalathur R, Liu L, Carlier PR. Structure of the G119S mutant acetylcholinesterase of the malaria vector Anopheles gambiae reveals basis of insecticide resistance. Structure. 2018 Jan 1;26(1):130.

14. Field LM, Emyr Davies TG, O’Reilly AO, Williamson MS, Wallace BA. Voltage-gated sodium channels as targets for pyrethroid insecticides. European Biophysics Journal. 2017 Oct 1;46(7):675.

15. Scott JG. Life and Death at the Voltage-Sensitive Sodium Channel: Evolution in Response to Insecticide Use. https://doi-org.ez.lshtm.ac.uk/101146/annurev-ento-011118-112420. 2019 Jan 10;64:243–57.

16. Kasai S, Itokawa K, Uemura N, Takaoka A, Furutani S, Maekawa Y, et al. Discovery of super– insecticide-resistant dengue mosquitoes in Asia: Threats of concomitant knockdown resistance mutations. Sci Adv. 2022 Dec 21;8(51).

17. Brengues C, Hawkes NJ, Chandre F, McCarroll L, Duchon S, Guillet P, et al. Pyrethroid and DDT cross-resistance in Aedes aegypti is correlated with novel mutations in the voltage-gated sodium channel gene. Med Vet Entomol. 2003 Mar;17(1):87–94.

18. Chung HH, Cheng IC, Chen YC, Lin C, Tomita T, Teng HJ. Voltage-gated sodium channel intron polymorphism and four mutations comprise six haplotypes in an Aedes aegypti population in Taiwan. PLoS Negl Trop Dis. 2019 Mar 1;13(3).

19. Fan Y, O’grady P, Yoshimizu M, Ponlawat A, Kaufmanid PE, Scott JG. Evidence for both sequential mutations and recombination in the evolution of kdr alleles in Aedes aegypti. PLoS Negl Trop Dis. 2020 Apr 1;14(4):1–22.

20. Weill M, Luffalla G, Mogensen K, Chandre F, Berthomieu A, Berticat C, et al. Insecticide resistance in mosquito vectors. Nature 2003 423:6936 [Internet]. 2003 May 8 [cited 2022 Aug 1];423(6936):136–7. Available from: https://www.nature.com/articles/423136b

21. Feyereisen R, Dermauw W, Van Leeuwen T. Genotype to phenotype, the molecular and physiological dimensions of resistance in arthropods. Pestic Biochem Physiol. 2015 Jun 1;121:61–77.

22. Engdahl C, Knutsson S, Fredriksson SÅ, Linusson A, Bucht G, Ekström F. Acetylcholinesterases from the Disease Vectors Aedes aegypti and Anopheles gambiae: Functional Characterization and Comparisons with Vertebrate Orthologues. PLoS One. 2015 Oct 8;10(10):e0138598.

23. Bloomquist JR. Toxicology, mode of action and target site-mediated resistance to insecticides acting on chloride channels. Comp Biochem Physiol C Pharmacol Toxicol Endocrinol. 1993 Oct 1;106(2):301–14.

24. Ffrench-Constant RH, Williamson MS, Davies TGE, Bass C. Ion channels as insecticide targets. 101080/0167706320161229781.2016 Oct 1;30(3–4):163–77.

25. Fonseca-González I, Quiñones ML, Lenhart A, Brogdon WG. Insecticide resistance status of Aedes aegypti (L.) from Colombia. Pest Manag Sci. 2011 Apr;67(4):430–7.

26. Goindin D, Delannay C, Gelasse A, Ramdini C, Gaude T, Faucon F, et al. Levels of insecticide resistance to deltamethrin, malathion, and temephos, and associated mechanisms in Aedes aegypti mosquitoes from the Guadeloupe and Saint Martin islands (French West Indies). Infect Dis Poverty. 2017 Feb 10;6(1).

27. Grau-Bové X, Tomlinson S, O’Reilly AO, Harding NJ, Miles A, Kwiatkowski D, et al. Evolution of the Insecticide Target Rdl in African Anopheles Is Driven by Interspecific and Interkaryotypic Introgression. Mol Biol Evol. 2020 Oct 1;37(10):2900–17.

28. Mintz JA. Two Cheers for Global POPs: A Summary and Assessment of the Stockholm Convention on Persistent Organic Pollutants. Georgetown International Environmental Law Review. 2001;14.

29. Wondji CS, Dabire RK, Tukur Z, Irving H, Djouaka R, Morgan JC. Identification and distribution of a GABA receptor mutation conferring dieldrin resistance in the malaria vector Anopheles funestus in Africa. Insect Biochem Mol Biol. 2011 Jul;41(7):484–91.

30. Yang C, Huang Z, Li M, Feng X, Qiu X. RDL mutations predict multiple insecticide resistance in Anopheles sinensis in Guangxi, China. Malar J. 2017 Nov 28;16(1).

31. Lumjuan N, Rajatileka S, Changsom D, Wicheer J, Leelapat P, Prapanthadara L aied, et al. The role of the Aedes aegypti Epsilon glutathione transferases in conferring resistance to DDT and pyrethroid insecticides. Insect Biochem Mol Biol. 2011 Mar;41(3):203–9.

32. Matthews BJ, Dudchenko O, Kingan SB, Koren S, Antoshechkin I, Crawford JE, et al. Improved reference genome of Aedes aegypti informs arbovirus vector control. Nature. 2018 Nov 22;563(7732):501–7.

33. Ortelli F, Rossiter LC, Vontas J, Ranson H, Hemingway J. Heterologous expression of four glutathione transferase genes genetically linked to a major insecticide-resistance locus from the malaria vector Anopheles gambiae. Biochem J. 2003 Aug 1;373(Pt 3):957–63.

34. Riveron JM, Yunta C, Ibrahim SS, Djouaka R, Irving H, Menze BD, et al. A single mutation in the GSTe2 gene allows tracking of metabolically based insecticide resistance in a major malaria vector. Genome Biol. 2014;15(2):R27.

35. Mitchell SN, Rigden DJ, Dowd AJ, Lu F, Wilding CS, Weetman D, et al. Metabolic and target-site mechanisms combine to confer strong DDT resistance in Anopheles gambiae. PLoS One. 2014 Mar 27;9(3).

36. Helvecio E, Romão TP, de Carvalho-Leandro D, de Oliveira IF, Cavalcanti AEHD, Reimer L, et al. Polymorphisms in GSTE2 is associated with temephos resistance in Aedes aegypti. Pestic Biochem Physiol. 2020 May 1;165.

37. Crava CM, Varghese FS, Pischedda E, Halbach R, Palatini U, Marconcini M, et al. Population genomics in the arboviral vector *Aedes aegypti* reveals the genomic architecture and evolution of endogenous viral elements. Mol Ecol. 2021 Jan 12;mec.15798.

38. Rose NH, Sylla M, Badolo A, Lutomiah J, Ayala D, Aribodor OB, et al. Climate and Urbanization Drive Mosquito Preference for Humans. Current Biology. 2020 Jul 23;

39. Lee Y, Schmidt H, Collier TC, Conner WR, Hanemaaijer MJ, Slatkin M, et al. Genome-wide divergence among invasive populations of Aedes aegypti in California. BMC Genomics. 2019 Mar 12;20(1):1–10.

40. Kelly ET, Mack LK, Campos M, Grippin C, Chen TY, Romero-Weaver AL, et al. Evidence of Local Extinction and Reintroduction of Aedes aegypti in Exeter, California. Frontiers in Tropical Diseases. 2021 Jul 8;2:703873.

41. Faucon F, Dusfour I, Gaude T, Navratil V, Boyer F, Chandre F, et al. Identifying genomic changes associated with insecticide resistance in the dengue mosquito Aedes aegypti by deep targeted sequencing. Genome Res. 2015 Sep 1;25(9):1347–59.

42. Leong CS, Vythilingam I, Liew JWK, Wong ML, Wan-Yusoff WS, Lau YL. Enzymatic and molecular characterization of insecticide resistance mechanisms in field populations of Aedes aegypti from Selangor, Malaysia. Parasit Vectors. 2019 May 16;12(1):1–17.

43. Poupardin R, Srisukontarat W, Yunta C, Ranson H. Identification of Carboxylesterase Genes Implicated in Temephos Resistance in the Dengue Vector Aedes aegypti. PLoS Negl Trop Dis. 2014;8(3).

44. Knox TB, Juma EO, Ochomo EO, Pates Jamet H, Ndungo L, Chege P, et al. An online tool for mapping insecticide resistance in major Anopheles vectors of human malaria parasites and review of resistance status for the Afrotropical region. Parasit Vectors. 2014 Feb 21;7(1):1– 14.

45. Langmead B, Salzberg SL. Fast gapped-read alignment with Bowtie 2. Nat Methods. 2012 Apr 4;9(4):357–9.

46. Li H, Handsaker B, Wysoker A, Fennell T, Ruan J, Homer N, et al. The Sequence Alignment/Map format and SAMtools. Bioinformatics. 2009 Aug 15;25(16):2078–9.

47. McKenna A, Hanna M, Banks E, Sivachenko A, Cibulskis K, Kernytsky A, et al. The Genome Analysis Toolkit: a MapReduce framework for analyzing next-generation DNA sequencing data. Genome Res. 2010 Sep;20(9):1297–303.

48. Danecek P, Bonfield JK, Liddle J, Marshall J, Ohan V, Pollard MO, et al. Twelve years of SAMtools and BCFtools. Gigascience. 2021 Feb 16;10(2).

49. Cingolani P, Platts A, Wang LL, Coon M, Nguyen T, Wang L, et al. A program for annotating and predicting the effects of single nucleotide polymorphisms, SnpEff: SNPs in the genome of Drosophila melanogaster strain w1118; iso-2; iso-3. Fly (Austin). 2012;6(2):80–92.

50. Boratyn GM, Schäffer AA, Agarwala R, Altschul SF, Lipman DJ, Madden TL. Domain enhanced lookup time accelerated BLAST. Biol Direct. 2012;

51. McInnes L, Healy J, Melville J. UMAP: Uniform Manifold Approximation and Projection for Dimension Reduction. ArXiv [Internet]. 2018 Feb 9 [cited 2021 Apr 22]; Available from: http://arxiv.org/abs/1802.03426

52. Campello RJGB, Moulavi D, Sander J. Density-Based Clustering Based on Hierarchical Density Estimates. Lecture Notes in Computer Science (including subseries Lecture Notes in Artificial Intelligence and Lecture Notes in Bioinformatics). 2013;7819 LNAI(PART 2):160–72.

53. Diaz-Papkovich A, Anderson-Trocmé L, Gravel S. A review of UMAP in population genetics. J Hum Genet. 2021 Jan 1;66(1):85–91.

54. Becht E, McInnes L, Healy J, Dutertre CA, Kwok IWH, Ng LG, et al. Dimensionality reduction for visualizing single-cell data using UMAP. Nat Biotechnol. 2018 Jan 1;37(1):38–47.

55. Bellin N, Calzolari M, Magoga G, Callegari E, Bonilauri P, Lelli D, et al. Unsupervised machine learning and geometric morphometrics as tools for the identification of inter and intraspecific variations in the Anopheles Maculipennis complex. Acta Trop. 2022 Sep 1;233:106585.

56. Evans R, O’Neill M, Pritzel A, Antropova N, Senior A, Green T, et al. Protein complex prediction with AlphaFold-Multimer. bioRxiv [Internet]. 2021 Oct 4 [cited 2022 Aug 2];2021.10.04.463034. Available from: https://www.biorxiv.org/content/10.1101/2021.10.04.463034v1

57. Jumper J, Evans R, Pritzel A, Green T, Figurnov M, Ronneberger O, et al. Highly accurate protein structure prediction with AlphaFold. Nature. 2021 Aug 26;596(7873):583–9.

58. Berman HM, Westbrook J, Feng Z, Gilliland G, Bhat TN, Weissig H, et al. The Protein Data Bank. Nucleic Acids Res. 2000 Jan 1;28(1):235–42.

59. Tatusova T, Dicuccio M, Badretdin A, Chetvernin V, Nawrocki EP, Zaslavsky L, et al. NCBI prokaryotic genome annotation pipeline. Nucleic Acids Res. 2016 Aug 19;44(14):6614–24.

60. Gloria-Soria A, Ayala D, Bheecarry A, Calderon-Arguedas O, Chadee DD, Chiappero M, et al. Global genetic diversity of Aedes aegypti. Mol Ecol. 2016 Nov 1;25(21):5377–95.

61. McBride CS, Baier F, Omondi AB, Spitzer SA, Lutomiah J, Sang R, et al. Evolution of mosquito preference for humans linked to an odorant receptor. Nature 2014 515:7526. 2014 Nov 12;515(7526):222–7.

62. Richly E, Leister D. NUMTs in sequenced eukaryotic genomes. Mol Biol Evol. 2004 Jun;21(6):1081–4.

63. Black Iv WC, Bernhardt SA. Abundant nuclear copies of mitochondrial origin (NUMTs) in the Aedes aegypti genome. Insect Mol Biol. 2009 Dec;18(6):705–13.

64. Clarkson CS, Miles A, Harding NJ, Lucas ER, Battey CJ, Amaya-Romero JE, et al. Genome variation and population structure among 1142 mosquitoes of the African malaria vector species Anopheles gambiae and Anopheles coluzzii. Genome Res. 2020 Oct 1;30(10):1533–46.

65. Colletier JP, Fournier D, Greenblatt HM, Stojan J, Sussman JL, Zaccai G, et al. Structural insights into substrate traffic and inhibition in acetylcholinesterase. EMBO J. 2006 Jun 21;25(12):2746–56.

66. Wang Y, Qiu L, Ranson H, Lumjuan N, Hemingway J, Setzer WN, et al. Structure of an insect epsilon class glutathione S-transferase from the malaria vector Anopheles gambiae provides an explanation for the high DDT-detoxifying activity. J Struct Biol. 2008 Nov 1;164(2):228–35.

67. Chen C, Compton A, Nikolouli K, Wang A, Aryan A, Sharma A, et al. Marker-assisted mapping enables forward genetic analysis in Aedes aegypti, an arboviral vector with vast recombination deserts. Genetics. 2022 Nov 1;222(3).

68. Hamid PH, Prastowo J, Widyasari A, Taubert A, Hermosilla C. Knockdown resistance (kdr) of the voltage-gated sodium channel gene of Aedes aegypti population in Denpasar, Bali, Indonesia. Parasit Vectors. 2017 Jun 5;10(1).

69. Rasli R, Lee HL, Ahmad NW, Fikri SFF, Ali R, Muhamed KA, et al. Susceptibility Status and Resistance Mechanisms in Permethrin-Selected, Laboratory Susceptible and Field-Collected Aedes aegypti from Malaysia. Insects 2018, Vol 9, Page 43. 2018 Apr 18;9(2):43.

70. Saha P, Chatterjee M, Ballav S, Chowdhury A, Basu N, Maji AK. Prevalence of kdr mutations and insecticide susceptibility among natural population of Aedes aegypti in West Bengal. PLoS One. 2019 Apr 1;14(4):e0215541.

71. Mano C, Jariyapan N, Sor-Suwan S, Roytrakul S, Kittisenachai S, Tippawangkosol P, et al. Protein expression in female salivary glands of pyrethroid-susceptible and resistant strains of Aedes aegypti mosquitoes. Parasit Vectors. 2019 Mar 14;12(1):1–19.

72. Yang F, Schildhauer S, Billeter SA, Yoshimizu MH, Payne R, Pakingan MJ, et al. Insecticide Resistance Status of Aedes aegypti (Diptera: Culicidae) in California by Biochemical Assays. J Med Entomol. 2020 Jul 1;57(4):1176.

73. Kandel Y, Vulcan J, Rodriguez SD, Moore E, Chung HN, Mitra S, et al. Widespread insecticide resistance in Aedes aegypti L. from New Mexico, U.S.A. PLoS One. 2019 Feb 1;14(2):e0212693.

74. Hernandez HM, Martinez FA, Vitek CJ. Insecticide Resistance in Aedes aegypti Varies Seasonally and Geographically in Texas/Mexico Border Cities. J Am Mosq Control Assoc. 2022 Mar 1;38(1):59–69.

75. Weill M, Luffalla G, Mogensen K, Chandre F, Berthomieu A, Berticat C, et al. Insecticide resistance in mosquito vectors. Nature 2003 423:6936. 2003 May 8;423(6936):136–7.

76. Chuaycharoensuk T, Juntarajumnong W, Boonyuan W, Bangs MJ, Akratanakul P, Thammapalo S, et al. Frequency of pyrethroid resistance in Aedes aegypti and Aedes albopictus (Diptera: Culicidae) in Thailand. Journal of Vector Ecology. 2011 Jun 1;36(1):204–12.

77. Solis-Santoyo F, Rodriguez AD, Penilla-Navarro RP, Sanchez D, Castillo-Vera A, Lopez-Solis AD, et al. Insecticide resistance in Aedes aegypti from Tapachula, Mexico: Spatial variation and response to historical insecticide use. PLoS Negl Trop Dis. 2021 Sep 1;15(9).

78. Ravenscroft TA, Janssens J, Lee PT, Tepe B, Marcogliese PC, Makhzami S, et al. Drosophila Voltage-Gated Sodium Channels Are Only Expressed in Active Neurons and Are Localized to Distal Axonal Initial Segment-like Domains. Journal of Neuroscience. 2020 Oct 14;40(42):7999–8024.

79. Vontas J, Katsavou E, Mavridis K. Cytochrome P450-based metabolic insecticide resistance in Anopheles and Aedes mosquito vectors: Muddying the waters. Pestic Biochem Physiol. 2020 Nov 1;170:104666.

80. Mugenzi LMJ, Menze BD, Tchouakui M, Wondji MJ, Irving H, Tchoupo M, et al. Cis-regulatory CYP6P9b P450 variants associated with loss of insecticide-treated bed net efficacy against Anopheles funestus. Nature Communications 2019 10:1. 2019 Oct 11;10(1):1–11.

81. Weedall GD, Mugenzi LMJ, Menze BD, Tchouakui M, Ibrahim SS, Amvongo-Adjia N, et al. A cytochrome P450 allele confers pyrethroid resistance on a major African malaria vector, reducing insecticide-treated bednet efficacy. Sci Transl Med. 2019 Mar 20;11(484):7386.

